# Dynamics of the CD9 interactome during bacterial infection of epithelial cells by proximity labelling proteomics

**DOI:** 10.1101/2024.12.13.628358

**Authors:** PA Wolverson, I Fernandes Parreira, MO Collins, JG Shaw, LR Green

**Affiliations:** School of Clinical Dentistry, University of Sheffield, Sheffield, United Kingdom; Florey Institute of Infection, University of Sheffield, Sheffield, United Kingdom; School of Biosciences, University of Sheffield, Sheffield, United Kingdom; School of Clinical Medicine and Population Health, University of Sheffield, Sheffield, United Kingdom

**Keywords:** Tetraspanin, CD9, proximity labelling, bacteria, *Neisseria meningitidis*, *Staphylococcus aureus*, epithelial

## Abstract

Epithelial colonisation is often a critical first step in bacterial pathogenesis, however, different bacterial species utilise several different receptors at the cell membrane to adhere to cells. We have previously demonstrated that interference of the human tetraspanin, CD9, can reduce adherence of multiple species of bacteria to epithelial cells by approximately 50%. However, CD9 does not act as a receptor and is responsible for organising and clustering partner proteins commandeered by bacteria for efficient adherence. CD9 can organise numerous host proteins at the cell membrane but the full interactome has not been delineated. Here, using a novel CD9 proximity-labelling model, we demonstrate a vast and diverse CD9 interactome with 845 significantly enriched proteins associated with CD9 over four hours. These putative proximal proteins were associated with various cellular pathways including cell adhesion, ECM-receptor interactions, endocytosis, SNARE interactions and adherens and tight junctions. Significant and known interactors of CD9 were enriched including β1 integrins and major immunoglobulin superfamily members but also included several known bacterial adherence receptors including CD44, CD46 and CD147. We further demonstrate dynamism of the interactome during infection at three separate time points with two different bacterial species, *Neisseria meningitidis* and *Staphylococcus aureus*. During meningococcal infection, 13 unique proximal proteins associated with CD9 were significantly enriched across four hours compared to uninfected cells. However, upon staphylococcal infection far fewer enriched proximal proteins were identified demonstrating that different bacteria require different host factors during CD9-mediated bacterial adherence. Transient knockdown of CD44 and CD147, candidate receptor proteins identified in our screen, significantly reduced staphylococcal and meningococcal adherence respectively. This effect was ablated in the absence of CD9 or if epithelial cells were treated with a CD9-derived peptide demonstrating the association of these proteins during staphylococcal and meningococcal adherence. We demonstrate for the first time the CD9 interactome of epithelial cells and that bacteria hijack these interactions to efficiently adhere to epithelial cells. This process is bacterial species specific, recruiting several different proteins during infection but a host-derived peptide is able to interfere with this process. We have therefore developed a tool that can measure changes within the CD9 interactome after cellular challenge, established a mechanism in which CD9 is used as a universal organiser of bacterial adhesion platforms and demonstrated that this process can be stopped using a CD9-derived peptide.

## Introduction

Tetraspanins are a superfamily of transmembrane proteins, consisting of 33 members in humans, that are classically characterised by four transmembrane domains (TM1-TM4), a small extracellular loop (EC1), a large extracellular loop (EC2), along with short intracellular N- and C-termini (1). Sequence and structural divergence within the extracellular loops and the intracellular domains, especially within the C-terminus, provides a unique function for each tetraspanin (2). A key characteristic of tetraspanins is their ability to associate with various membrane proteins, including integrins, proteoglycans and immunoglobulin superfamily members (3, 4), to form molecular complexes known as tetraspanin-enriched microdomains (TEMs) (5). TEMs are involved in myriad biological processes, including cell adhesion, migration and signalling, and have been identified as a route of infection for many viral and bacterial pathogens (5, 6).

Recent studies have implicated tetraspanins in various bacterial adherence pathways, including those of pathogenic *Neisseria* spp. (7), *Staphylococcus aureus* (4, 8), *Escherichia coli* (9), *Burkholderia pseudomallei* (10, 11) and *Mycobacterium abscessus* (12). In most cases, the tetraspanins themselves do not act as the bacterial receptor but coordinate organisation and clustering of receptors that allow for bacterial adherence, suggesting the composition of the TEM is important for initial bacterial attachment (4, 7). For example, CD9 promotes *S. aureus* adherence to epithelial cells through organisation of the syndecans, allowing recruitment of fibronectin to create an optimal adhesive platform for the bacterium (4). However, each bacterial species that exploits tetraspanin-mediated adherence co-opt different receptors during cellular invasion but the tetraspanins have been demonstrated to associate with many of these membrane proteins.

*Neisseria meningitidis*, an opportunistic bacterial pathogen able to cause invasive meningococcal disease (IMD), uses various bacterial adhesins to adhere to many different cell types (13). The canonical adherence pathway consists of an initial interaction between type IV pili and either CD147 (14) or potentially CD46 (15). From here, an intimate association occurs after interaction of the opacity proteins (Opa) with either heparan sulphate proteoglycans (HSPGs) or CEACAMs (16). Similarly, *S. aureus*, a common human commensal which can also cause multiple infectious diseases including pneumonia, osteomyelitis and infective endocarditis, utilises a plethora of adhesins to associate with multiple host cell receptors (17). Perhaps the most studied of these is the interaction of the staphylococcal fibronectin binding proteins to fibronectin which is utilised as a ‘bridge’ with α5β1 integrin (18). Still, several other pathways have been investigated and include co-opting host receptors such as CD36, desmoglein 1 and annexin 2 (17). The tetraspanin CD9 has previously been demonstrated to associate with various integrins (19), HSPGs (4), immunoglobulin superfamily members (3), CD36 (20) and CD46 (21) in the formation of TEMs.

Here, we utilise proximity labelling to characterise the CD9 interactome on epithelial cells and demonstrate that different CD9 associated TEMs are required for infection by different bacteria. We also report dynamism of these TEMs during infection with specific proteins recruited to CD9 associated TEMs over time. Finally, we demonstrate that a CD9-derived peptide can disrupt these interactions and reduce specific bacterial adherence to cells. Disruption of TEMs and therefore disruption of the formation of bacterial cell surface receptor complexes is arising as a promising therapeutic target and could aid in overcoming the growing pressures associated with antimicrobial resistance.

## Materials and Methods

### Strains and bacterial growth conditions

The *N. meningitidis* and *S. aureus* strains used in this study were serogroup B MC58 and SH1000 respectively. *N. meningitidis* solid cultures were grown on brain:heart infusion (BHI; Oxoid, Ltd., UK) agar with 10% Levinthal’s solution (22) overnight at 37°C with 5% CO2, while staphylococcal strains were grown on Luria broth (LB; Oxoid) agar overnight at 37°C. Meningococcal liquid cultures were grown in BHI broth with 5% Levinthal’s solution and 10 mM sodium bicarbonate at 37°C with constant agitation. SH1000 was grown in LB at 37°C with constant agitation. All liquid cultures were inoculated using freshly grown plates.

### Cell culture

Wild-type (WT) and CD9 knockout (CD9^-/-^) A549 human lung epithelial cells (a kind gift from David Blake (Fort Lewis College, Colorado, USA) (23)) were maintained in Dulbecco’s Modified Eagle’s Media (DMEM; Thermo Fisher Scientific, Massachusetts, USA) and 10% heat-inactivated fetal calf serum (FCS; Bioscience, UK).

### Peptides and antibodies

The CD9 EC2-derived peptide (800C; DEPQRETLKAIHYALN) and scrambled peptide (Scr; QEALKYNRAETPLDIH) were designed as described previously (4) and synthesized using solid phase Fmoc chemistry (Genscript, New Jersey, USA). The peptide, 800C, has been demonstrated to inhibit staphylococcal interactions with human cells without affecting bacterial growth (4).

Mouse anti-human CD9 IgG1 (MM2/57; Merck, Germany), mouse anti-human CD9 IgG1 (602.29; kind gift of Lynda Partridge, University of Sheffield, Sheffield, UK), mouse anti-human CD147 IgG1 (HIM6; Biolegend, California, USA), mouse anti-human CD46 IgG1 (MEM-258; Thermo Fisher Scientific), mouse anti-human CEACAM1 IgG1 (B3-17; Merck), mouse anti-human CEACAM6 IgG1 (1H7-4B; Merck), mouse anti-human CD44 IgG2a (60224-1-Ig; Proteintech Group, Inc, Illinois, USA), mouse anti-human GAPDH (MAB374; Merck), mouse IgG1 (JC1; in house), mouse IgG2a (02-6200; Thermofisher Scientific), mouse anti-FLAG (M2; Merck), goat anti-mouse HRP (P0447; Agilent, California, USA) were used as described.

### Flow cytometry

Flow cytometry was used to measure tetraspanin and bacterial receptor expression. Cell dissociation buffer (Thermo Fisher Scientific) was used to detach adherent cells before labelling with the relevant antibody at 4°C for 60 minutes. If required, cells were secondary labelled with a fluorescein isothiocyanate (FITC) conjugated goat anti-mouse IgG antibody (F5897, Merck). Cells were fixed with 1% paraformaldehyde, quantified with an LSRII cytometer (Becton Dickinson, Oxford, UK) and analysed by FlowJo v10.0.7r2 software (BD).

### Western blots

Cell lysates were prepared in RIPA buffer (10mM Tris-HCl pH8.0, 140mM NaCl, 0.5mM EGTA, 1mM EDTA, 0.1% Sodium Deoxycholate, 0.1% SDS, 1% Triton X-100) with a protease inhibitor cocktail (Complete Mini, Roche, Switzerland) and fractionated on SDS-PAGE gels. Fractionated proteins were blotted on to a nitrocellulose membrane, blocked with non-fat dried milk diluted in TBST, and subsequently probed with the appropriate antibody and visualized using an ECL detection system (Merck). To blot for biotinylated protein, nitrocellulose blots were blocked in 5% BSA and probed with a streptavidin HRP conjugate (Thermo Fisher Scientific).

### Plasmids

The lentiviral vector used to express CD9-TurboID in our study was constructed and packaged by Vectorbuilder (VB220615-1144dtm; Vectorbuilder Inc., Illinois, USA). The vector was designed to fuse TurboID to human CD9 at the C-terminus separated by a 3 tandem GGGGS linker under the control of a CMV promoter. A V5 tag was added to the C terminus of TurboID. An eGFP lentiviral control vector (VB010000-9389rbj; Vectorbuilder) was used throughout the study. Addition of the puromycin resistance gene allowed for selection and maintenance of cells containing the construct. Lentiviral transfection was carried out as previously described (4). Recombinant lentivirus was produced by co-transfecting sub-confluent HEK293T cells with the pMD2G (Addgene, Massachusetts, USA), psPAX2 (Addgene) and pLV[Exp]-CD9-TurboID or the pLV[Exp]-eGFP vectors with jetPEI (Polyplus transfection, Illkirch-Graffenstaden, France). Media was changed after 24 hours and lentivirus-containing supernatants were harvested after 48 hours. For stable transfection, WT or CD9^-/-^ cells were grown for 24 hours, 20% of culture media was replaced with lentivirus-containing supernatant. Polybrene was used to increase transduction efficiency. Transduction was checked by protein electrophoresis and Western blotting.

### Fluorescence microscopy

CD9 was visualized as described previously (7). Cells were grown overnight on glass coverslips and fixed with 1% paraformaldehyde. Coverslips were washed, blocked in 5% goat serum (Merck) and treated with anti-CD9 antibody followed by goat anti-mouse FITC conjugated antibody to visualize CD9. Cells were washed and stained with DAPI (4′,6-diamidino-2-phenylindole) to visualize cell nuclei. Coverslips were mounted with Vectashield mounting medium with DAPI (Vector Labs, California, USA), allowing visualization on a Leica DMRB upright fluorescent microscope.

### Calcein AM adhesion assay

Black tissue-culture treated 96 well plates were coated with 75µg/ml fibronectin (Merck) for three hours. Wells were washed with PBS and seeded with 2x10^4^ cells for one hour or 24 hours. Cells were washed twice with PBS and stained with 0.01µM Calcein AM for one hour. Fibronectin-coated wells without cells were used as a blank control. Fluorescence was read using the FLUOstar-OPTIMA plate reader (488/520nm). Cell adherence was calculated as a percentage of adherence by WT cells, set at 100%.

### Infection assays

Infection assays were carried out as previously described (4). Cells were seeded onto 96 well plates and cultured overnight. To reduce non-specific binding, wells were blocked with 5% bovine serum albumin (BSA; Merck). Cells were treated with peptide for 60 minutes before infection with either MC58 or SH1000 at a multiplicity of infection (MOI) of 50 for 60 minutes at 37°C with 5% CO2. Cells were washed and lysed with 2% saponin (Merck) for 30 minutes. Serial dilutions of lysates were plated onto BHI with 10% Levinthal’s solution or LB agar plates and allowed to grow overnight. The number of bacteria bound to BSA blocked empty wells was subtracted from adherent and internalized bacteria. Bacterial adherence to peptide treated cells was calculated as a percentage of bacterial adherence to untreated cells, set at 100%.

To analyse the CD9 interactome during infection, CD9^-/-^ or CD9^-/-^ cells transfected with CD9-TurboID were grown overnight to confluency in 100mm tissue culture dishes. Dishes were blocked for 60 minutes with 5% BSA (Merck) before infection with MC58 or SH1000 at an MOI of 50. Uninfected and infected cells were treated with 50µM biotin at the point of infection. Infection was allowed to proceed for 30, 60 or 240 minutes before cells were washed, harvested with cell dissociation solution (Thermo Fisher Scientific) and lysed in cold RIPA buffer.

### Biotin pull-down

Biotinylated proteins were extracted from cell lysates using Dynabeads^TM^ M-280 streptavidin magnetic beads (Thermo Fisher Scientific). 50µl of beads were washed in cold RIPA buffer without protease inhibitors before cell lysates were added and incubated with end over end rotation, overnight at 4°C. Supernatants were removed and consecutively washed with the following; i) 2% SDS/50mM Tris pH7.4, ii) cold RIPA buffer, iii) 2M urea/50mM ammonium bicarbonate, iv) 50mM ammonium bicarbonate.

### Mass spectrometry

Streptavidin purifications were reduced and alkylated using 5mM tri-(2-carboxyethyl) phosphine hydrochloride (TCEP) and 10mM iodoacetamide respectively. 2µg trypsin (MS Grade; Pierce, Thermo Fisher Scientific) was added to each sample for on-bead digestion and incubated for 3 hours at 37°C. Peptides were acidified through addition of trifluoroacetic acid (TFA) and desalted using C18 columns (Pierce, Thermo Fisher Scientific) and eluted peptides were dried using a vacuum concentrator (Eppendorf) at 45°C. Peptides were resuspended in 12 μL 0.5% formic acid and 5 μL of each sample was analyzed by nanoflow LC-MS/MS using an Orbitrap Exploris 480 (Thermo Fisher) mass spectrometer with an EASY-Spray source coupled to a Vanquish LC System (Thermo Fisher). Peptides were desalted online using a Pepmap Neo C18 nano trap column, 300 μm I.D.X 5 mm (Thermo Fisher) and then separated using an EASY-Spray column, 50 cm × 75 μm ID, PepMap Neo C18, 2 μm particles, 10 Å pore size (Thermo Fisher). The 90 min gradient was used, starting from 3% to 20% buffer B (0.5% formic acid in 80% acetonitrile) for 60 min, then ramping up to 35% buffer B for 15 mins, then up to 99% buffer B for 1 min and maintaining at 99% buffer B for 9 min. The Orbitrap Exploris was operated in positive mode with a DDA cycle time of 2 seconds. MS1 spectra were acquired at a resolution of 120,000 at m/z 200, with a scan range (m/z) 375–1,200, the standard AGC target. The most abundant multiply charged (2+ and higher) ions in a given chromatographic window were subjected to HCD fragmentation with a collision energy of 30% and dynamic exclusion set to automatic. The MS2 AGC target was set to standard, and MS2 spectra were measured with a resolution setting of 30,000 at m/z 200.

### Proteomic data analysis

All raw mass spectrometry data were processed with MaxQuant version 1.6.10.43. Data were searched against a human (July 2022), *S. aureus* NCTC 8325 (May 2023) and *N. meningitidis* MC58 (July 2023) UniProt sequence database using the following search parameters: digestion set to Trypsin/P with a maximum of 2 missed cleavages, methionine oxidation and N-terminal protein acetylation as variable modifications, cysteine carbamidomethylation as a fixed modification, match between runs enabled with a match time window of 0.7 min and a 20 min alignment time window, label-free quantification enabled with a minimum ratio count of 2, minimum number of neighbours of 3 and an average number of neighbours of 6. A first search precursor tolerance of 20 ppm and a main search precursor tolerance of 4.5 ppm was used for FTMS scans and a 0.5 Da tolerance for ITMS scans. A protein FDR of 0.01 and a peptide FDR of 0.01 were used for identification level cut-offs. MaxQuant output was loaded into Perseus version 1.6.10.50 and the matrix was filtered to remove all proteins that were potential contaminants, only identified by site and reverse sequences. LFQ intensities were log2(x) transformed and data was filtered to retain proteins with a minimum of three valid LFQ Intensities in one group. Subsequently, data were visualised using multi-scatter plots and Pearson’s correlation analysis. Data were normalised by subtracting column medians and missing values were imputed from the normal distribution with a width of 0.3 and downshift of 1.8. To identify quantitatively enriched proteins between groups, two-sided Student’s t-tests were performed with a permutation-based FDR of 0.05 with an *S*0 = 0.1. The mass spectrometry proteomics data have been deposited to the ProteomeXchange Consortium via the PRIDE (24) partner repository with the dataset identifier PXD058283. Data was exported into an Excel file and input into GraphPad Prism to create the figures and plots presented. Gene ontology and pathway analysis was performed using SubcellulaRVis (25) and WebGestalt (26).

### siRNA knockdown

Target genes were knocked down using siGENOME SMARTpool human CD44 siRNA (M-009999-03), human CD46 siRNA (M-004570-00) or human CD147 siRNA (M-010737-01).

Non-targeting siRNA pool (D-001206-13; Horizon Discovery Ltd., Cambridge, UK) was used as an appropriate control. 1.5 x 10^5^ WT or CD9^-/-^ cells were seeded onto 6 well plates and cultured overnight. 5 µL of siRNA was mixed with 245 µL of serum free media, while 3 µL of Dharmafect 1 (Horizon Discovery) was added to 247 µL of serum free media. Both were left at room temperature for 5 minutes before mixing and incubated for a further 20 minutes before adding dropwise to cells. Cells were incubated for 48 hours before harvesting with trypsin/EDTA and preparing for infection assays in 96 well plates as described. Knockdown was confirmed by flow cytometry.

### Statistical analyses

All analyses were performed within GraphPad Prism version 10.2.2 (GraphPad Software Inc., USA). Significance was established at *p*≤0.05, all data represents at least three independent experiments. Statistical considerations and specific analyses are described separately within each section. * specify significance to the untreated control unless otherwise specified; * *p*≤0.05, ** *p*≤0.01, *** *p*≤0.001.

## Results

### Disruption of CD9 can affect both meningococcal and staphylococcal adherence to epithelial cells

Tetraspanins have previously been demonstrated to be involved in both meningococcal and staphylococcal adherence (4, 7). These data were confirmed by measuring meningococcal and staphylococcal adherence to wild-type (WT) A549 or tetraspanin CD9^-/-^ A549 cells. Knockout of CD9 significantly reduced both meningococcal (58.7±6.1%) and staphylococcal (49.2±8.7%) attachment compared to WT cells (Fig. 1A). Furthermore, a CD9-derived peptide, 800C, can also affect both bacterial adherence pathways. Pre-treatment of WT cells with 800C significantly reduced meningococcal adherence to cells compared to untreated cells (56.5±8.8%; Fig. 1B). Similarly, as previously demonstrated (4), 800C treatment of WT cells significantly reduced staphylococcal adherence to cells in comparison to untreated cells (41.0±9.9%; Fig. 1C). A scrambled 800C peptide control had no effect on either bacterial adherence pathway. Adherence of both bacteria was significantly reduced in CD9^-/-^ cells, however, treatment with the scrambled peptide or 800C had no further effect (Fig. 1B-C).

**Figure 1.**
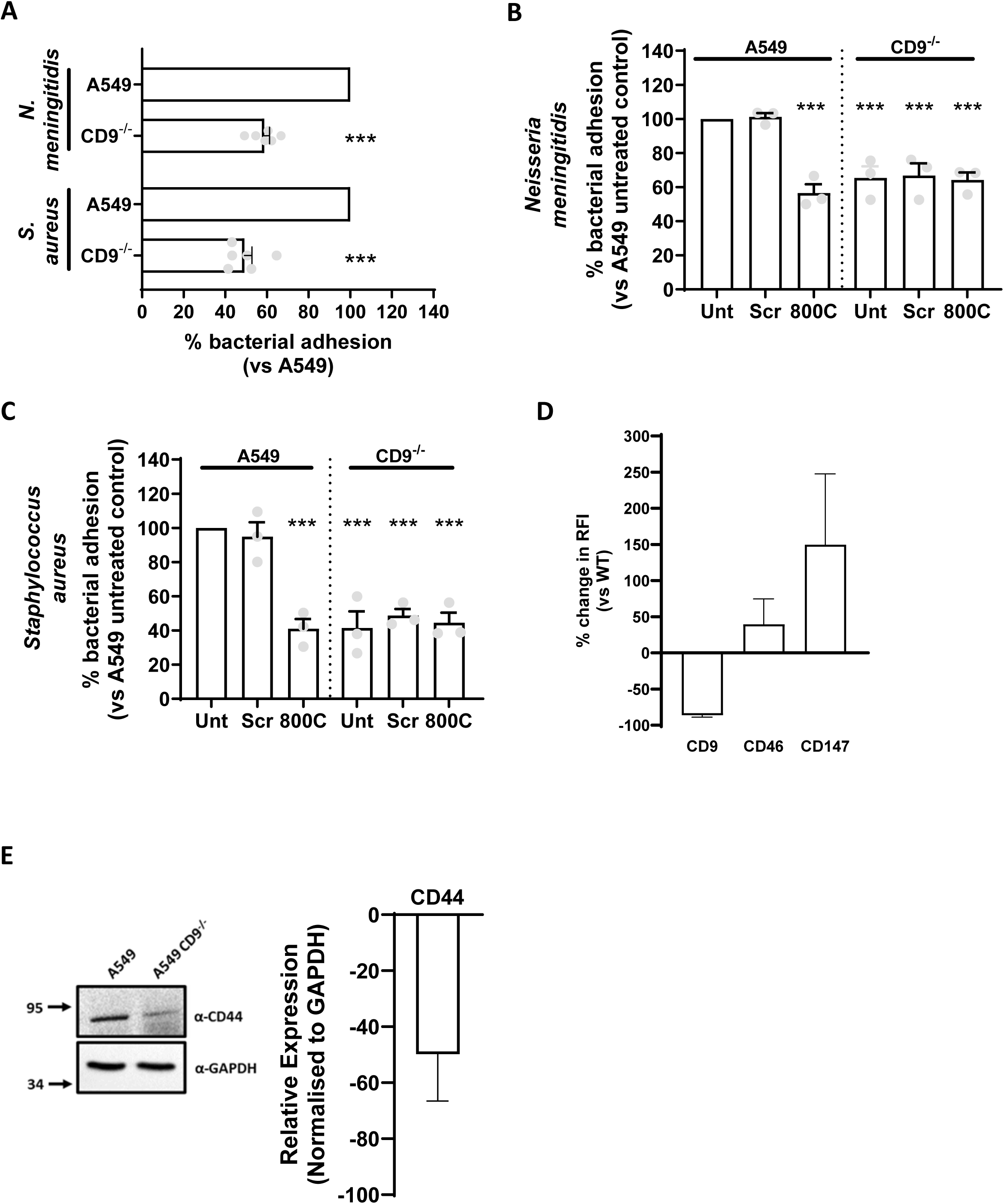
CD9 knockout affects both meningococcal and staphylococcal adherence and changes receptor expression. (A) CD9 knockout disrupts both meningococcal and staphylococcal adherence. WT and CD9^-/-^ cells were infected with either meningococci (MC58) or staphylococci (SH1000) for 60 mins at an MOI=50. Cells were disrupted after infection and adherent and internalised bacteria were enumerated by colony forming units (cfu). (B-C) CD9-derived peptide, 800C, reduces meningococcal adherence (B) and staphylcoccal adherence (C) to epithelial cells. WT or CD9^-/-^ cells were pre-treated with a scrambled peptide or 800C for 60 mins prior to infection. Cells were infected as above and adherence enumerated by cfu. (D) Cell surface expression of meningococcal receptors measured by flow cytometry. WT or CD9^-/-^ cells were treated with an anti-CD9 antibody (602.29), an anti-CD46 antibody, an anti-CD147 antibody. A mouse IgG isotype control (JC1) was included and expression was determined using a FITC-conjugated secondary antibody. % change was calculated by comparing WT to CD9^-/-^ cells. (E) Representative blot demonstrating expression of meningococcal and staphylococcal receptors. Whole cell lysates of WT and CD9^-/-^ cells were electrophoresed and blotted. Blots were probed with an anti-CD44 antibody. An anti-GAPDH antibody was used as a loading control. Densitometry was calculated through ImageJ analysis, removing background with an empty lane and normalising to the loading control. n≥3, mean ± SEM.

CD9 expression was substantially reduced in CD9^-/-^ cells in comparison to the WT cells (-86.33±4.4%; Fig. 1D). Previously, we have observed that the staphylococcal receptor, syndecan-1, is affected by removal of CD9 (4). Here, we investigate the expression of known meningococcal receptors (CD46, CD147, CEACAM1 and CEACAM6) and a known glycoprotein implicated in staphylococcal adherence (CD44) to epithelial cells (27). Analysis of cells by flow cytometry demonstrated that removal of CD9 increased the expression of CD46 (39.64±35.0%) and CD147 (149.7±97.9%) at the cell membrane, however, these increases were not significant (Fig. 1B). No significant differences were observed in the percentage of cells expressing our target proteins (Supp. Fig. 1A). Conversely, decreases were observed by Western blot in CD44 expression after CD9 removal (-49.8±16.8%). Expression of CEACAM1 and CEACAM6 was also reduced after CD9 removal (Supp. Fig. 1B). Thus, we have confirmed that removal of tetraspanin CD9 can affect multiple bacterial adherence pathways despite changes in expression in putative receptors.

### Efficient labelling of CD9 with TurboID

We have demonstrated that CD9 interference can affect very distinct bacterial adherence pathways. We have also previously shown that CD9 does not appear to act as a receptor for these bacteria (7), placing the impetus for tetraspanin-mediated adherence on their proximal proteins. *N. meningitidis* and *S. aureus* utilise different canonical receptors for adhesion suggesting that either differing CD9 platforms are required or that specific proteins are recruited to CD9 during adherence. To investigate this, we employed a proximity labelling methodology, fusing CD9 to the promiscuous biotin ligase, TurboID, which allows rapid labelling (1-10 min) and identification of proximal proteins (∼10 nm) (28). TurboID was fused to the C-terminus of CD9 in line with previous strategies which have tagged CD9 (29). A flexible linker was added to ensure access to the C-terminus and allow free movement of TurboID (Fig. 2A). The construct also contains a V5-tag at the TurboID C-terminus as another method of detecting the protein. Stable transfection of A549 WT and CD9^-/-^ cells followed by Western blotting revealed that the construct was expressed, with both endogenous CD9 and the CD9 fusion present in WT cells (Fig. 2B). Expression levels of the CD9:TurboID fusion protein were similar to those of the endogenous protein. Efficient biotinylation of a range of proteins at different molecular weights was observed after 10 minutes of addition of biotin to the media of transfected cells in comparison to WT and CD9^-/-^ cells (Fig. 2C). Continued accumulation of biotinylated proteins was observed between 1 to 18 hours. Biotinylated proteins were efficiently pulled down using streptavidin beads from CD9^-/-^ transfected cells in comparison to CD9^-/-^ cells (Fig. 2D). Therefore, we have developed a model by which we can measure the proximal proteins of CD9 over time and with differing stimuli.

**Figure 2.**
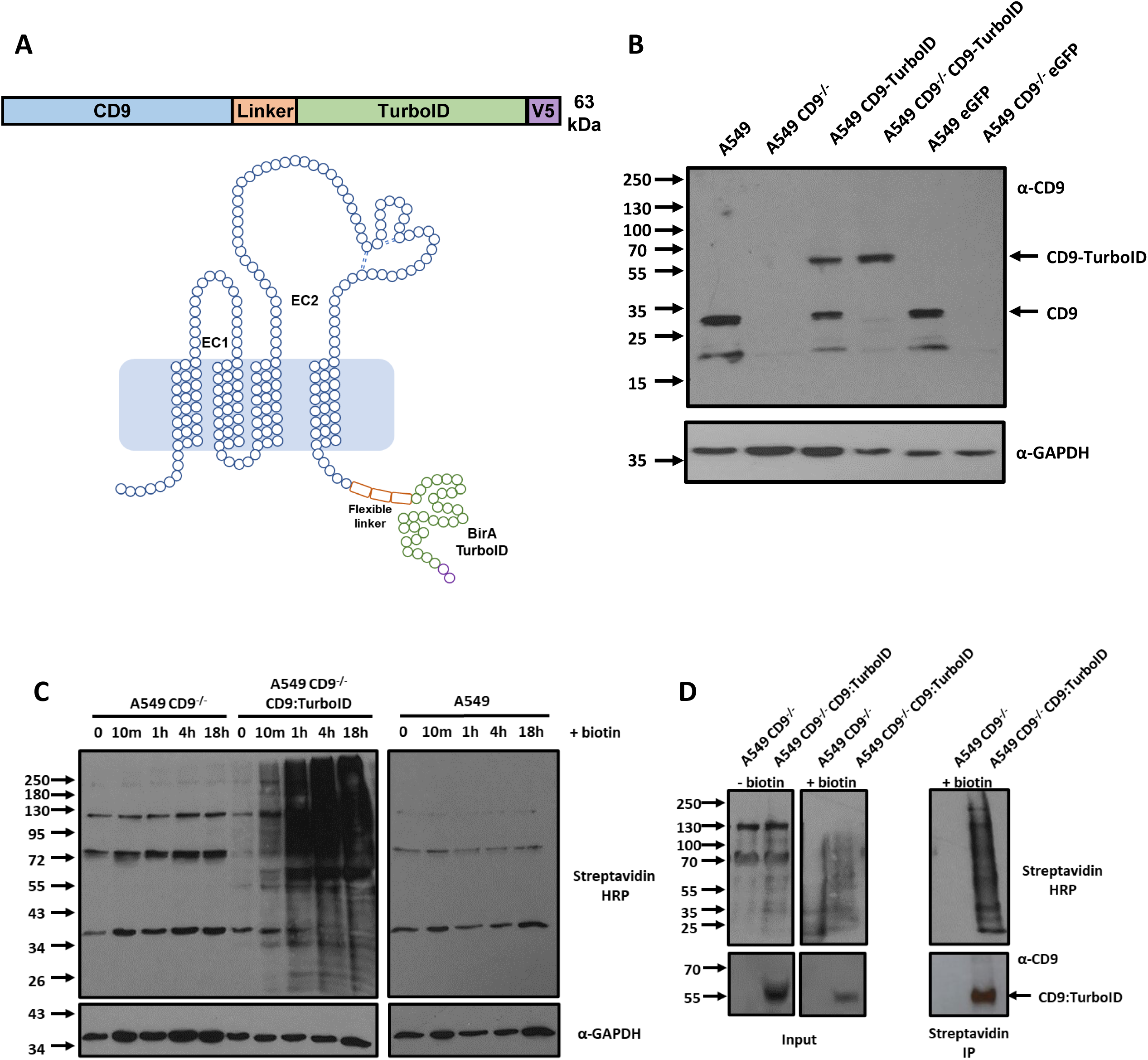
CD9:TurboID fusion construct is expressed and localises to the plasma membrane. (A) Schematic of the CD9:TurboID fusion protein used in this study. The relative position of a flexible linker, TurboID and a V5 tag are indicated. (B) Whole cell lysates of WT and CD9^-/-^ cells and those transfected with either the CD9:TurboID fusion plasmid or the eGFP control plasmid were electrophoresed and blotted. Blots were probed with anti-CD9 (MM2-59) and anti-GAPDH antibodies. (C) Representative blot demonstrating biotinylated proteins produced by WT, CD9^-/-^ or CD9^-/-^ CD9:TurboID transfected cells. Cells were treated with 50µM biotin for 18 hours and sampled at various timepoints. Blotted proteins were probed with a streptavidin HRP conjugate. (D) Streptavidin bead pull-downs demonstrate a large number of enriched biotinylated proteins. CD9^-/-^ or transfected cells were treated with 50µM biotin for 60 minutes. Biotinylated proteins were pulled down from lysed cells using streptavidin beads. The input and resulting pull-down were probed with a streptavidin HRP conjugate.

### CD9:TurboID fusion protein functions similar to endogenous CD9

To ensure that the CD9:TurboID fusion protein can function similarly to endogenous CD9, as alteration of the C-terminus may affect normal function (30), we investigated both the localisation and biological activity of CD9 during bacterial infection. CD9 was abundantly expressed on the plasma membrane of WT cells in comparison to CD9^-/-^ or isotype control treated cells (Fig. 3A). Stably transfected CD9^-/-^ cells demonstrated a recovery of CD9 expression at the cell surface with both transfected cells and WT cells exhibiting strong expression at intercellular junctions with non-punctate expression patterns. Calcein AM assays were used to test the ability of cells to adhere to fibronectin over time. CD9^-/-^ cells demonstrated significantly reduced adherence to fibronectin over one hour (50.0±13.4%) compared to WT cells, however, this phenotype was partially rescued through stable transfection with the CD9:TurboID fusion protein (80.9±11.0%; Fig. 3B). No significant difference was observed in cell adhesion to fibronectin after 24 hours. Epithelial cells were also checked for their ability to accommodate bacterial infection, as we have previously demonstrated that CD9 is important for multiple adherence pathways (Fig. 1). As expected, both meningococcal (57.7±3.5%) and staphylococcal (44.1±1.5%) adherence was significantly reduced in the CD9^-/-^ cells compared to WT and WT cells transfected with an eGFP construct (Fig. 3C). This was mirrored in CD9^-/-^ cells transfected with an eGFP construct. However, in CD9^-/-^ cells transfected with the CD9:TurboID construct, both meningococcal (90.0±2.6%) and staphylococcal (85.6±8.6%) adherence increased similar to levels observed in WT or eGFP transfected WT cells (Fig. 3C). Interestingly, despite overexpression of CD9 in CD9:TurboID transfected cells, only a small increase was observed in both meningococcal (107.1±6.4%) or staphylococcal (114.4+12.8%) adherence. We therefore demonstrate that CD9 can function normally despite fusion of TurboID to the C-terminus.

**Figure 3.**
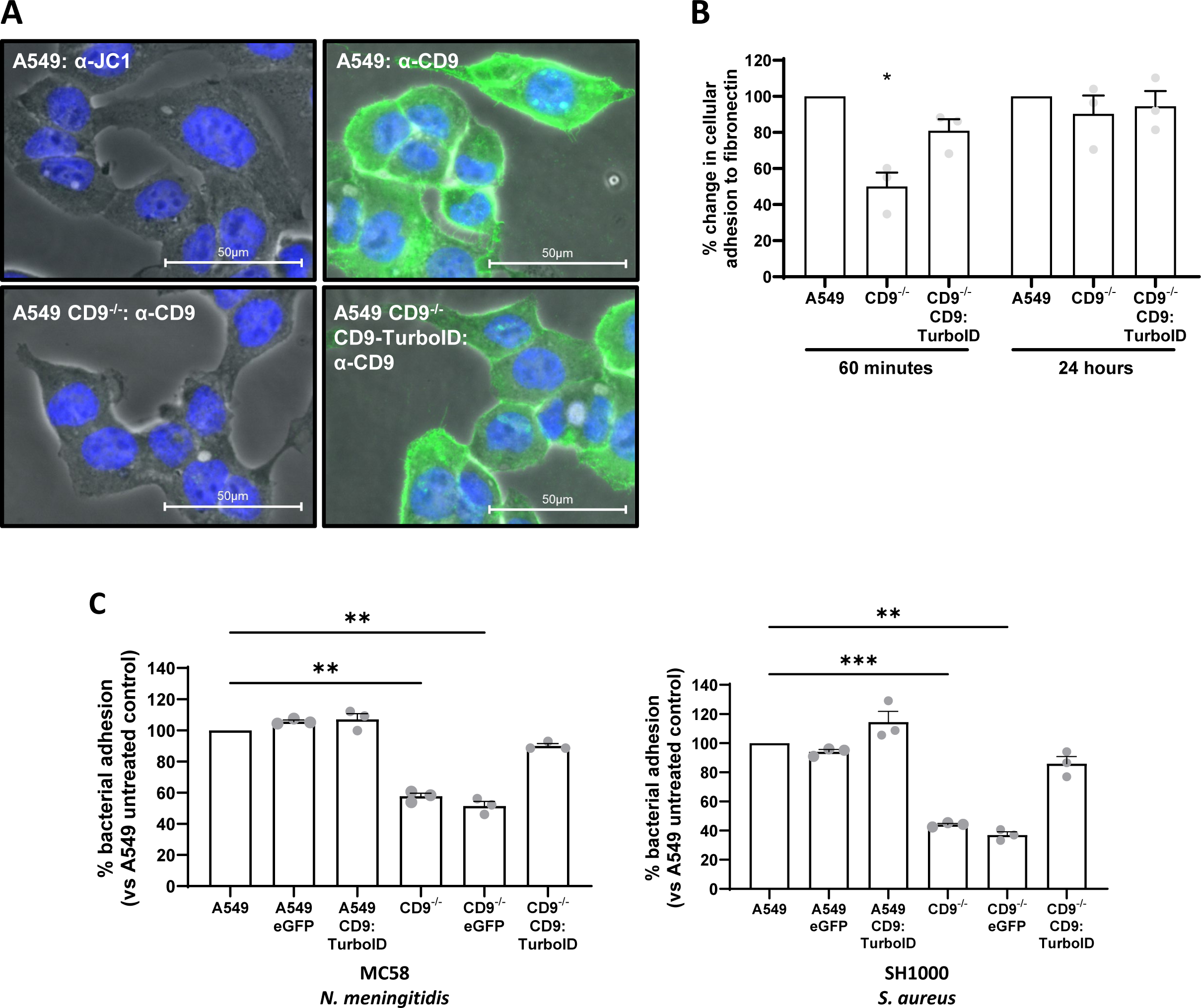
CD9:TurboID biotinylates proteins and rescues a meningococcal and staphylococcal infection phenotype. (A) Representative immunofluorescence images showing localisation of CD9 constructs. WT and CD9^-/-^ cells and those transfected with the CD9:TurboID fusion plasmid were fixed and treated with anti-CD9 (602.29) antibodies. Localisation was visualised using a FITC-conjugated secondary antibody. (B) The cell adhesion phenotype of CD9 is rescued with expression of the CD9:TurboID fusion protein in CD9^-/-^ cells. Calcein AM labelled cells were allowed to adhere to fibronectin coated wells for 60 minutes or 24 hours. (C) Meningococcal and staphylococcal infection phenotypes are rescued with expression of CD9:TurboID fusion protein in CD9^-/-^ cells. Cells were infected with meningococci (MC58) or staphylococci (SH1000) for 60 minutes at an MOI=50. Cells were disrupted after infection and adherent and internalised bacteria were enumerated by cfu. n>3, mean ± SEM, One-Way ANOVA.

**TurboID identifies several known CD9 interactors and significantly extends the number of putative proximal proteins**

To generate a definitive picture of the CD9 interactome we purified biotinylated proteins over three time points, 30, 60 and 240 minutes, to capture proximal proteins involved with the proposed myriad functions of CD9. The volcano plots demonstrate the individual isolated proteins enriched through biotinylation by CD9:TurboID (Fig. 4A). 845 proteins were isolated across all three time points with 351, 633 and 821 proteins enriched at 30, 60 and 240 minutes respectively (Fig. 4A-B). 231 enriched proteins were unique across timepoints (30 mins – 0; 60 mins – 21; 240 mins – 210), 268 were shared between two timepoints and 346 proteins were shared across all three timepoints suggestive of a core interactome (Fig. 4B). The most enriched proteins were discoidin (DCBCLD1), an integral membrane protein, at 30 minutes, an <u>a</u>dhesion <u>m</u>olecule with <u>Ig</u> like domain 2 (AMIGO2), at 60 minutes and TACC2, a cytoplasmic protein associated with centrosomes during the cell cycle, at 240 minutes. Several known CD9 interactors were identified including CD9 itself, several integrins (α5 and β1), other tetraspanin members (CD151, Tspan15) and immunoglobulin superfamily members including IGSF3 (Fig. 4A, Supp. Data 1). Furthermore, several putative meningococcal and staphylococcal receptors were identified including CD147, CD46, CD44 and syndecan-1 (Fig. 4A), which we have previously demonstrated directly interacts with CD9 (4). Interestingly, CD46 is not within the enriched cohort of proximal proteins at 30 minutes, lying just outside of the stringency cutoff (s0=1), but is abundant at both 60 and 240 minutes. Despite changes in CEACAM expression after CD9 knockout (Supp. Fig. 1B), no CEACAMs were observed in the enriched proximal protein dataset.

**Figure 4.**
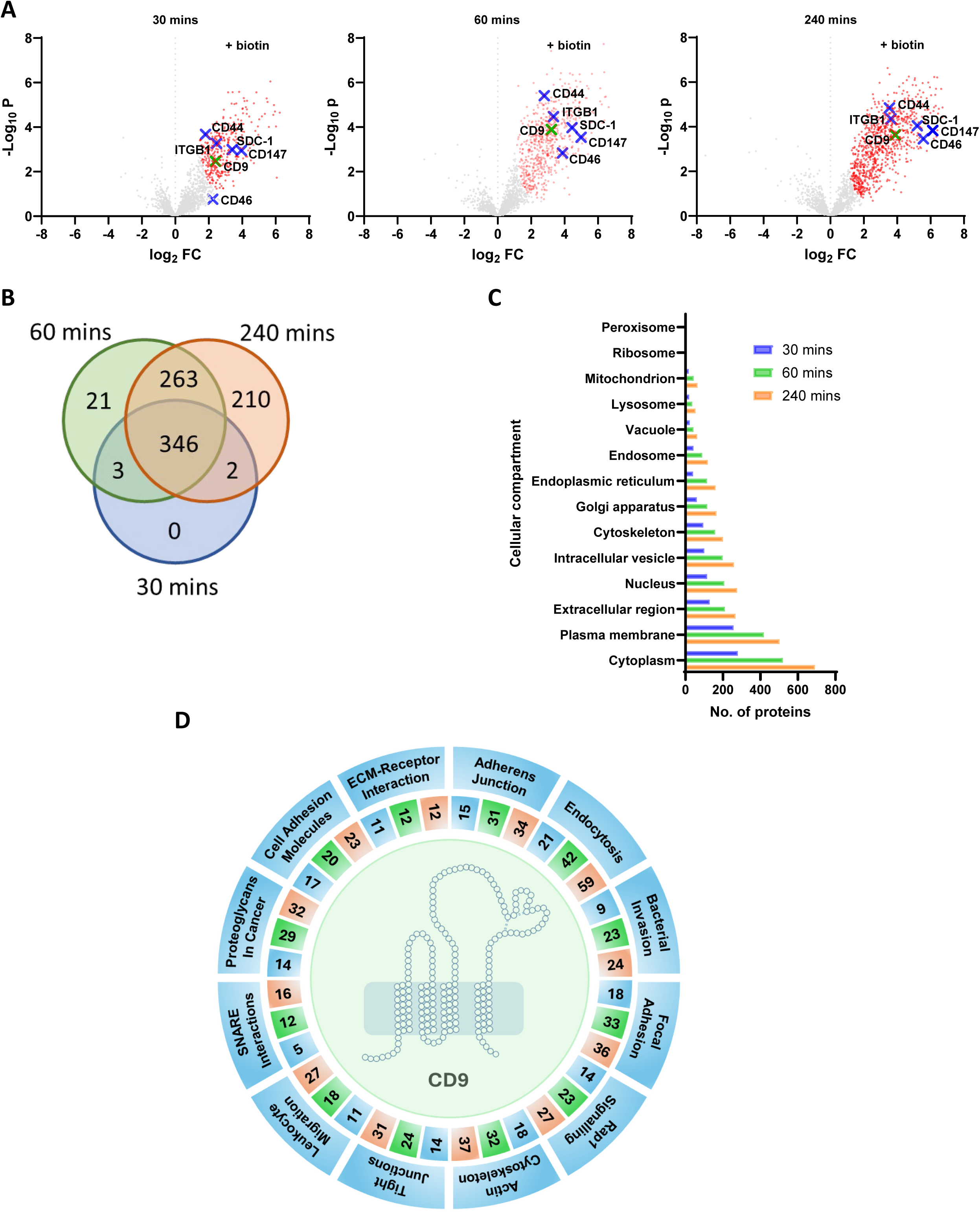
The CD9 interactome is large and encompasses several proteins involved in myriad cell functions. (A) Volcano plots of LFQ data from CD9^-/-^ cells transfected with the CD9:TurboID fusion protein versus WT cells treated with 50µM biotin over time. Significantly enriched proteins (red) for the CD9:TurboID fusion protein were determined by Student’s t-test at stringency (s0=1). Known meningococcal and staphylococcal receptors are shown (blue crosses) as is CD9 (green cross). (B) Venn diagram demonstrating the overlap and dynamism of the CD9 interactome over time. Significantly enriched proteins were compared between timepoints, 30 mins blue; 60 mins green; 240 mins orange. (C) The majority of proximal proteins are found within the cytoplasm and the plasma membrane. Significantly enriched proteins over time were analysed through SubcellulaRVis and categorised by their cellular compartments. (D) Selected biological functions associated with CD9 proximal proteins. Significantly enriched proteins were analysed by KEGG pathway. Selected enriched KEGG pathways (FDR ≤ 0.05) are shown. Numbers demonstrate proximal proteins associated with the pathway by time point, 30 mins blue; 60 mins green; 240 mins orange.

Through analysis of the enriched CD9 interactome by cellular compartment, as expected most proteins observed were found in either the plasma membrane or the cytoplasm (240 mins – 502 and 691 respectively) but a large number were also found in intracellular vesicles (240 mins – 259; Fig. 4C). Many proteins were also associated with the cytoskeleton (240 mins – 200), which supports the role of CD9 in cell adhesion and migration but is also suggestive of our study isolating various signalling proteins associated with these processes. Interestingly, we also observed several proximal proteins associated with the extracellular region (240 mins – 267; Fig. 4C). In all cases, the number of proximal proteins associated with each cellular compartment rose over time. Cellular localisation of the enriched proximal proteins demonstrates successful fusion and tagging of CD9.

Using a KEGG pathway analysis, we identified CD9 involvement in several cellular processes (Fig. 4D). Using a false discovery rate less than or equal to 0.05, 24 pathways were identified after 30 minutes, 39 after 60 minutes and 46 after 240 minutes (Supp. Data 2). Across all timepoints, several of these pathways have previously been linked with CD9 including adherens junctions, tight junctions, endocytosis, and cell adhesion molecules (31) (Fig. 4D). Other expected pathways, such as SNARE interactions, were also observed (30 mins, 5/33; 60 mins, 12/33; 240 mins, 16/33), with numbers of proteins involved with these pathways increasing over time. Despite no infectious challenge, proteins associated with several pathways involved in bacterial or viral infection were identified across all time points (30 mins, 4; 60 mins, 8; 240 mins, 10), with specific correlations to *E. coli* invasion, *Yersinia* infection, Shigellosis and *Salmonella* infection (Supp. Data 2). Numerous pathways often requisitioned by bacteria during infection were also identified including regulation of the actin cytoskeleton, critical in actin remodelling at the plasma membrane during infection, and ECM-receptor interaction, which includes important groups of proteins often commandeered for use during bacterial adherence. The latter contained several integrins and proteoglycans, previously identified as important during bacterial adherence, such as α5β1, CD44 and syndecan-1 (4, 18, 27), but also other unidentified proteins such as dystroglycan-1 which require further investigation. Interestingly, several signalling pathways were also identified during our proximity labelling screen, in particular, proteins associated with Rap1 signalling, a GTPase which modulates expression and activation of integrins and matrix metalloproteases (32), were observed throughout each timepoint (30 mins, 14/210; 60 mins, 23/210; 240 mins, 27/210; Fig. 4D, Supp. Data 2). Therefore, using proximity labelling we have described the CD9 interactome, delineating several known proteins and novel CD9 interactors whilst simultaneously identifying a role for CD9 in both canonical and non-canonical pathways.

### The CD9 interactome is dynamic and changes with bacterial infection

As we have observed that numerous proteins associated with bacterial adherence pathways are enriched during our CD9 proximity labelling screen and that interference with CD9 can diminish bacterial adherence, we have investigated whether the CD9 interactome can change during meningococcal or staphylococcal adherence. The volcano plots demonstrate that thirteen proteins proximal to CD9 are enriched during meningococcal infection in comparison to uninfected cells (30 mins, 4; 60mins, 2; 240 mins 7; Fig. 5A). The most enriched proteins associated with meningococcal infection were YTHDF3 at 30 minutes, NNMT at 60 minutes and CISD1 at 240 minutes. Conversely, three CD9 proximal proteins were enriched in uninfected cells compared to cells infected with meningococcal bacteria (60 mins, 2; 240 mins, 1). In contrast, only one protein that associates with CD9 was enriched during staphylococcal infection after 60 minutes, PRPF40A (Fig. 5A). All proteins identified at each timepoint were unique to the specific infecting bacteria. Specific functions for the identified proteins are provided (Table 1).

**Figure 5.**
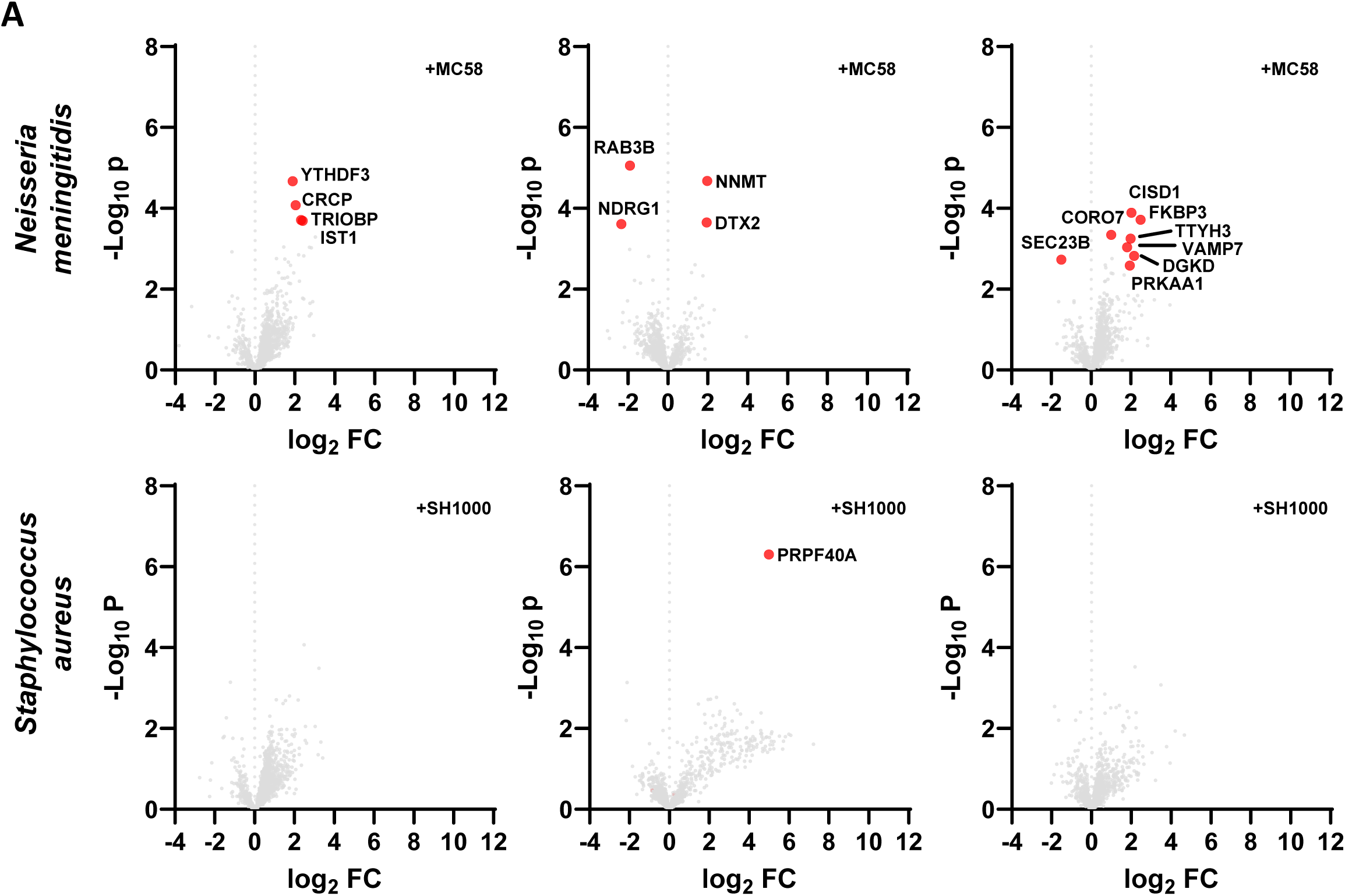
The CD9 interactome is dynamic and changes dependent on infection. (A) Volcano plots of LFQ data from infected CD9^-/-^ cells transfected with the CD9:TurboID fusion protein versus uninfected CD9^-/-^ cells transfected with the CD9:TurboID fusion protein. Cells were treated with 50µM biotin and infected with either meningococci (MC58) or staphylococci (SH1000) at an MOI=50 for the stated time period. Significantly enriched proteins (red) were determined by Student’s t-test at stringency (s0=0.1).

**Table 1.** Functions and cellular process of significantly enriched CD9 proximal proteins observed during infection. Table shows both proteins enriched in infected (orange) vs uninfected cells (blue) for both meningococci and staphylococci (green).

| Gene | Full name | Function | Classification |  |
| --- | --- | --- | --- | --- |
| YTHDF3 | YTH N6-methyladenosine RNA binding protein F3 | Binding of RNA | mRNA biogenesis | 30 minutes |
| CRCP | CGRP receptor component | Calcitonin receptor increases cAMP levels | Transcription machinery |  |
| TRIOBP | TRIO and F-actin binding protein | Interacts with TRIO, stabilises F-actin | Cytoskeletal proteins |  |
| IST1 | IST1 factor associated with ESCRT-III | Interacts with endosomal sorting complexes | Endocytosis |  |
| Gene | Full name | Function | Classification |  |
| RAB3B | RAB3B, member RAS oncogene family | GTPase activity. Regulation of vesicle size | Membrane trafficking | 60 minutes |
| NDRG1 | N-myc downstream regulated 1 | Stress responses, p53-mediated caspase activation | Exosome |  |
| NNMT | Nicotinamide N-methyltransferase | Nicotinamide metabolism | Metabolism of cofactors |  |
| DTX2 | Deltex E3 ubiquitin ligase 2 | E3 ubiquitin ligase | Notch signaling pathway |  |
| PRPF40A | Pre-mRNA processing factor 40 homolog A | Involved in cytoskeleton organisation | Transcription |  |
| Gene | Full name | Function | Classification |  |
| SEC23B | SEC23 homolog B, COPII coat complex component | Vesicle trafficking | Membrane trafficking | 240 minutes |
| CORO7 | Coronin 7 | F-actin regulator | Cytoskeleton proteins |  |
| CISD1 | CDSGH iron sulphur domain 1 | Regulation of oxidation | Transport |  |
| FKBP3 | Prolyl isomerase 3 | Role in immunoregulation, protein folding and trafficking | Chaperones |  |
| TTYH3 | Tweety family member 3 | Chloride anion channels | Transporters |  |
| VAMP7 | Vesicle-associated membrane protein 7 | Fusion of transport vesicles to target membranes | SNARE interactions |  |
| DGKD | Diacylglycerol kinase | Phosphorylates diacylglycerol to produce phosphatidic acid | PI signalling pathway |  |
| PRKAA1 | Protein kinase AMP-activated subunit alpha 1 | Ser/Thr protein kinase | Tight junctions |  |

Several of these proximal proteins are involved in typically appropriated pathways during infection including cytoskeletal rearrangement, membrane trafficking, transport and signalling. 11/17 proteins were uniquely enriched at their specific timepoint, however, YTHDF3, TRIOBP, IST1 and DTX2 were enriched in meningococcal infected cells at 30 or 60 minutes, but also in untreated cells at 240 minutes, suggesting meningococcal infection induces recruitment to sites proximal to CD9 earlier than usual. TRIOBP is involved in cytoskeletal reorganisation and protein trafficking suggesting an earlier recruitment could facilitate meningococcal induced membrane perturbation (33). YTHDF3 has been linked to immunosuppression signalling responses during infection (34), similar to another candidate also recruited at 30 minutes, CRCP (35), and suggests CD9 involvement in dampening the meningococcal inflammatory response.

Interestingly, RAB3B and NDRG1, enriched in untreated cells compared to those with meningococcal infection at 60 minutes, were also found in both infected and uninfected cells at 240 minutes, suggesting meningococcal infection delays their recruitment. Rab proteins have a role in endocytosis and meningococci have previously been found to co-localise in the subapical region of Rab11b, Rab22a and Rab3 positive vesicles (36). Whereas, low NDRG1 expression has previously been reported in damaged epithelial cells and associated with tight junction disruption (37).

At 240 minutes post meningococcal infection, several proteins involved in vacuole trafficking (VAMP7, CORO7), DNA damage responses (FKBP3, DTX2), inflammation and immune cell activation (TTYH3, DGKD, PRKAA1) were enriched compared to the uninfected control. Interestingly, presence of the SNARE family member, VAMP7, can increase bacterial growth within intracellular vacuoles and therefore enhance intracellular survival (38). Finally, we observed a reduction in enrichment of SEC23B after meningococcal infection for 240 minutes, intriguingly, enteropathogenic *Escherichia coli* has previously been shown to disrupt the Sec23/Sec24 complex and disrupt host protein transport as a virulence mechanism (39). Overall, we demonstrate that the CD9 interactome can change during bacterial infection, but these changes are dependent on the infectious agent.

### The CD9 interactome includes known meningococcal and staphylococcal receptors which are involved in CD9-mediated bacterial adherence

We specifically demonstrate enrichment of several known meningococcal and staphylococcal receptors amongst the CD9 interactome. As we and others have previously demonstrated direct CD9 interaction of β1 integrins and syndecan-1 in staphylococcal adhesion (4, 40), two important hits from our screen, we wanted to further demonstrate that CD9 and other receptor candidates from the interactome were involved in tetraspanin-mediated bacterial adherence and that this was dependent on the infecting bacteria. We therefore focused on CD46 and CD147 as putative pilus receptors of the meningococcus and CD44 as a known interactor in various bacterial adherence pathways including *Escherichia coli* (41), Group A Streptococcus (GAS) (42) and *S. aureus* (27). Using pooled siRNA we demonstrated successful knockdown of each of our target proteins (Fig. 6A). After siRNA-mediated knockdown of these proteins we observed significant reductions in meningococcal adherence to CD147-depleted epithelial cells (56.4±3.1%) compared to untreated or non-targeting siRNA treated cells (Fig. 6B). CD46 or CD44 knockdown did not impact meningococcal adherence to epithelial cells suggesting the importance of CD147, but not CD46 or CD44. As observed previously, addition of a CD9-derived peptide (800C) to untreated and non-targeting siRNA treated cells significantly reduced meningococcal adherence (45.7±3.0% and 47.8±3.6% respectively) but no effect was demonstrated after treatment with a scrambled control peptide (Fig. 6B). This significant reduction was also observed in CD46- and CD44-depleted cells. However, no significant reduction was observed between CD147-depleted cells with no peptide treatment and CD147-depleted cells treated with 800C. This suggests no major additive effect of these treatments and that CD147 and CD9 interact within the same pathway during meningococcal adherence. As before, meningococcal adherence to CD9^-/-^ cells was significantly reduced compared to the wild-type cells (51.1±5.3%; Fig. 6B). However, as with all siRNA treatments and peptide treatments, CD147-depletion had no effect on meningococcal adherence in CD9^-/-^ cells (Fig. 6B) further suggesting the importance of interaction between these two proteins during tetraspanin-mediated adherence.

**Figure 6.**
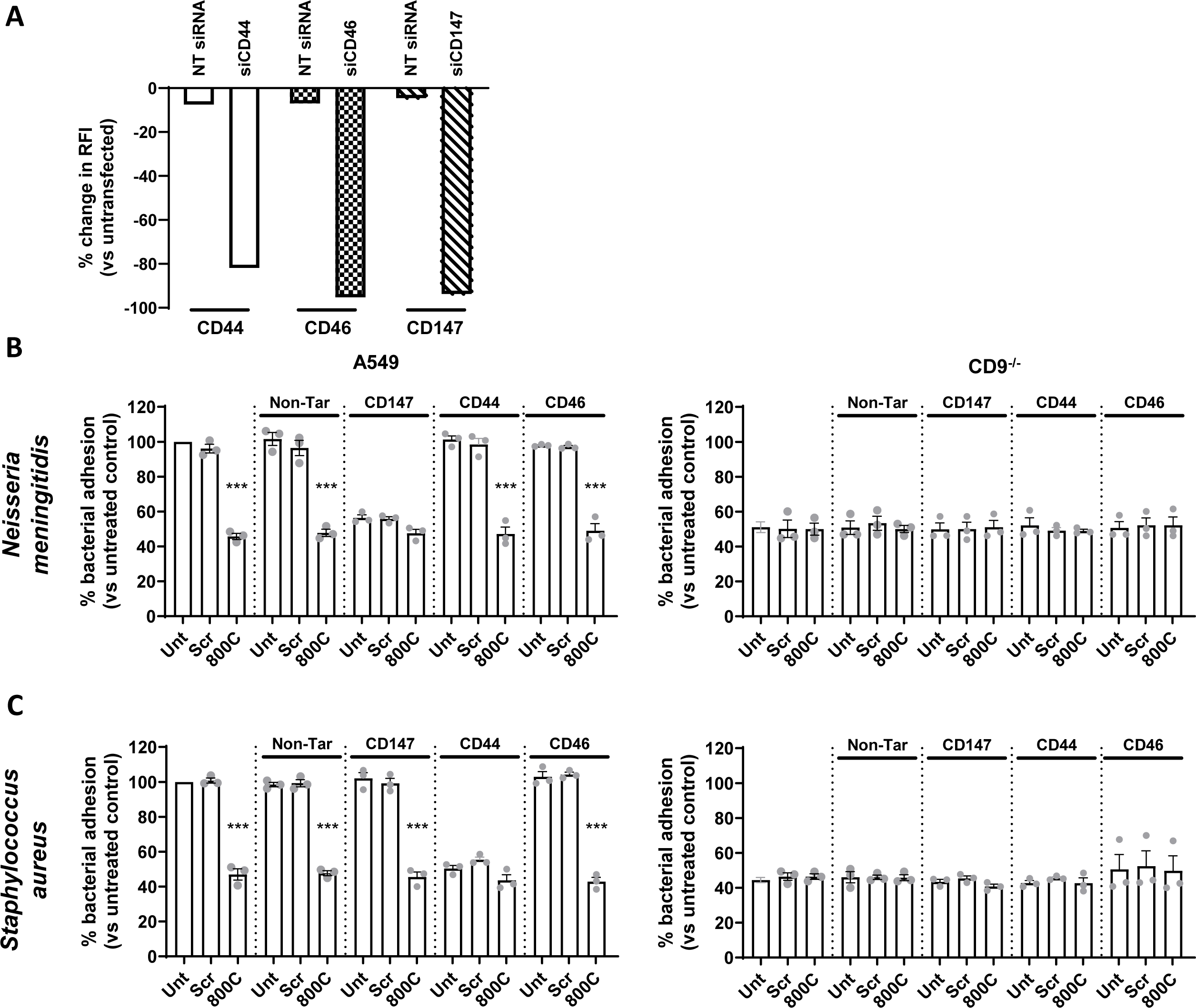
Differing CD9 proximal proteins are involved in CD9-mediated meningococcal and staphylococcal adhesion. WT cells were treated with specific siRNAs 72 hours prior to infection. Cells were treated with 800C or a scrambled control peptide for 60 minutes prior to infection. Cells were infected with either meningococci (MC58) or staphylococci (SH1000) for 60 mins at an MOI=50. (A) Flow cytometry demonstrating efficient knockdown of meningococcal and staphylococcal receptors after siRNA treatment prior to infection. Cells were probed with anti-CD147, anti-CD46 and anti-CD44 antibodies. n=1, mean. (B) CD147 knockdown and CD9-derived peptide treatment demonstrate no additive effects on meningococcal adherence. WT and CD9^-/-^ siRNA treated cells were infected with meningococci (MC58) for 60 mins at MOI=50. (C) CD44 knockdown and CD9-derived peptide treatment demonstrate reduced additive effects in staphylococcal adherence. WT and CD9^-/-^ siRNA treated cells were infected with staphylococci (SH1000) for 60 mins at MOI=50. Cells were disrupted after infection and adherent and internalised bacteria were enumerated by cfu. n=3, mean ± SEM, One-Way ANOVA.

Similarly, significant reductions were observed in staphylococcal adherence to CD44-depleted cells (50.4±3.0%) compared with untreated and non-targeting siRNA treated cells (Fig. 6C). However, no effects were demonstrated in CD46- or CD147-depleted epithelial cells suggesting CD44 is important during staphylococcal adherence. Upon addition of 800C, staphylococcal adherence was significantly reduced in untreated cells (47.0±5.6%), non-targeting siRNA cells (47.8±2.4%), CD46- (42.9±4.2%) and CD147-depleted cells (45.6±4.9%; Fig. 6C). Addition of a scrambled control peptide had no effect on any of the siRNA-treated cells. However, no significant reduction in adherence was observed when CD44-depleted cells with no peptide were compared to CD44-depleted cells treated with 800C. As before, this suggests that the interaction between CD44 and CD9 is important during staphylococcal adherence. Staphylococcal adherence was significantly reduced in CD9^-/-^ cells compared to wild-type cells but no further reductions were observed with any siRNA or peptide treatments, including CD44-depletion (Fig. 6C). These data further suggest the importance of an interaction between CD9 and CD44 during tetraspanin-mediated staphylococcal adherence. In conclusion, we have demonstrated that known meningococcal and staphylococcal receptors, identified within our CD9 proximal protein screen, are important during tetraspanin-mediated bacterial adherence and that, critically, these proximal proteins differ between the infecting organism but CD9 remains as a universal constant during bacterial adherence.

## Discussion

In this study, we have characterised the CD9 interactome using a proximity labelling approach with the promiscuous biotin ligase, TurboID, fused to CD9. We demonstrate that fusion of TurboID to the C-terminus of CD9 has no effect on CD9 localisation and function, and use of the fusion protein reestablishes bacterial adherence to cells lacking CD9. We have identified several novel and known enriched proximal proteins associated with CD9. Many of these proximal proteins are involved in mammalian cell adhesion and their associated signalling pathways, however, there is also a prevalence for proteins utilised during bacterial adherence and invasion pathways. We demonstrate enrichment and recruitment of specific proteins during meningococcal adherence but not during staphylococcal adherence, suggesting dynamic and mobile CD9 microdomains during tetraspanin-mediated bacterial adherence. Finally, we demonstrate involvement of CD147 and CD44 in CD9-mediated meningococcal and staphylococcal adherence respectively, but not vice versa confirming the requirement for specificity of proximal proteins during tetraspanin-mediated bacterial adherence.

Tetraspanins have been observed to interact several partner proteins (43), however, these studies have often focused on the interaction of the tetraspanin with a specific partner. Various techniques have been used to populate a partial interactome for CD9 and the other tetraspanins including immunoprecipitation (4), super resolution microscopy (44, 45), single molecule tracking (46), cryo-EM (47), yeast two-hybrid assays (48) and affinity purification (49). However, these techniques often require knowledge of potential tetraspanin interactors, require high levels of expression of the target proteins, struggle to evaluate weak or transient interactions and provide limitations to the screen size. Recently, attempts have been made to broadly define the tetraspanin interactome using proximity labelling techniques (50). Proximity labelling using TurboID followed by mass spectrometry allows rapid biotinylation of proximal proteins to the target protein, providing a broad interactome over time and after specific challenges to the cell which also considers weak or transient interactions (28). We have developed a tool to measure the CD9 interactome in epithelial cells by fusing TurboID to the C-terminus of CD9.

We believe this is the first study of the broader CD9 interactome by proximity labelling, identifying 845 potential proximal proteins after incubation with exogenous biotin for 4 hours. This included known interactors such as IGSF3 (51), HSPGs (4), various integrins (19), metalloproteases (52) and other tetraspanins but also several novel interactors involved in signal transduction, transportation and cell migration. Similarly, Cheerathodi *et al.* identified 1,600 proteins within the CD63 interactome after 36 hours with enrichment of Rab GTPases and SNARE proteins (50). Whilst we observed similar pathways and proteins we also describe enrichment of many tight and adherens junction proteins, perhaps unsurprising considering the canonical localisation of CD9 to the plasma membrane in comparison to CD63 localisation within the lysosome. Surprisingly, we detected enrichment of several extracellular proteins, however, we suspect that a cohort of these proteins are not included within our interactome due to fusion of TurboID to the cytoplasmic tail of CD9. For example, despite the robust presence of cell membrane surface HSPGs, such as syndecan-1 and CD44, within our screen we did not detect any glycosylphosphatidylinositol (GPI)-anchored proteins, such as the glypicans or specific members of the CEACAM family. However, the lack of enrichment of any CEACAM members, either GPI-anchored or otherwise, suggests that CD9 does not directly associate with these proteins but they may be indirectly associated with the tetraspanin through more transient interactions within the cell membrane. Future studies could focus on CD9-interacting extracellular proteins by utilising new proximity labelling techniques (53). However, CD9 enrichment of both membrane proteins and effectors associated with cell signalling cascades provides further evidence of the role of CD9 as a coordinator of the plasma membrane but also downstream proteins and cellular processes.

We have previously demonstrated that CD9 does not act as a bacterial receptor and that their partner proteins are required for efficient bacterial adherence (4, 7), similar to viral utilisation of the tetraspanins (54). Here, we confirmed, through transient knockdown, that two highly enriched CD9 proximal proteins, CD147 and CD44, were associated with CD9-mediated meningococcal and staphylococcal adherence respectively. CD147 is a known meningococcal receptor (14) whilst CD44 has been demonstrated as a phagocytic receptor of *S. aureus* in macrophages (27) and was recently enriched during a *S. aureus* proximity labelling screen during endothelial cell infection (55). This, alongside our previous data that two other significantly enriched proximal proteins, α5β1 integrin and syndecan-1, are involved in CD9-mediated staphylococcal adherence (4), suggests that proximity labelling of CD9 proximal proteins enriches numerous candidate proteins during bacterial infection. However, we observed no change in meningococcal adherence after transient knockdown of CD46 suggesting no involvement of this putative receptor (15) in CD9-mediated bacterial adherence despite enrichment of the protein at both 60 and 240 minutes in uninfected cells.

Similar to viral remodelling of TEMs (50), we observed dynamism in CD9-enriched microdomains, however, enrichment of cognate partner proteins was aligned with infection by specific bacteria. At all timepoints, infection with meningococci led to more enriched CD9 proximal receptors and downstream effectors than infection with staphylococci. We hypothesise that the required receptors for staphylococcal infection, particularly those required for bacterial adhesion after 30 minutes, are likely within pre-formed CD9-enriched microdomains, and thus its recruitment is less dynamic than during meningococcal infection. For example, we observed high enrichment of both α5β1 integrin and syndecan-1 within uninfected cells, which are linked to canonical and non-canonical staphylococcal adherence pathways (4, 18). Although, we note that certain proteins enriched during meningococcal infection (DTX2, DGKD and VAMP7) were also present during staphylococcal infection but just outside the significance threshold. Meningococci possess a robust phase variation mechanism allowing rapid switching of expression states for many virulence factors critical for bacterial adherence allowing association with various host surface proteins (56). Therefore, in the absence of a known effector molecule released by the bacteria to signal for membrane composition changes, and the speed of bacterial adherence within our model, changes in bacterial outer membrane virulence factors could cause shifts within CD9-enriched microdomains over time. Future studies could utilise more targeted proximity labelling techniques, such as the use of APEX or APEXII (57). This may delineate further dynamics within CD9-enriched microdomains as biotinylation begins upon addition of hydrogen peroxide therefore allowing specific timepoints and protein localisation to be addressed, however, the addition of hydrogen peroxide may have a detrimental effect on the bacteria.

Our study also enriched proteins involved in pathways utilised by both bacteria and viruses to enter cells and provided similar partner proteins to those previously observed in other tetraspanin studies. For example, during HPV entry to cells CD63 interaction with ALIX, an ESCRT-associated adaptor protein, and syntenin-1, a syndecan binding protein, is critical (54). Interestingly, during meningococcal adherence we observed early enrichment of IST-1, an ESCRT-associated protein, suggesting that membrane remodelling and endocytosis are important for infection. Similarly, the potential recruitment of VAMP7 during both staphylococcal and meningococcal adherence would suggest the involvement of endocytic pathways and survival within the cell, which has been reported within both pathogens (58, 59). Furthermore, the enrichment of CD9 proximal proteins involved in cytoskeletal regulation and vesicular transport in this study would suggest traversal and survival within the cell. Conversely, the CD9 interactome also contained several tight and adherens junction proteins. Bacterial induced remodelling of the cell surface, particularly of basolateral proteins involved in tight junctions can promote bacterial traversal, particularly of the blood:brain barrier (60). We observed delayed enrichment of NDRG1, with low expression of this protein associated with disruption of tight junctions. The involvement of CD9 proximal proteins in both putative intracellular and paracellular traversal routes and in membrane and cytoskeleton remodelling suggests multiple mechanisms by which bacteria could manipulate tetraspanins for adherence and may explain their involvement in many bacterial infections (7, 9, 11).

We have successfully generated a tool to track the CD9 interactome in epithelial cells using proximity labelling and demonstrated dynamism within these microdomains in response to infection with either *N. meningitidis* or *S. aureus*. We demonstrate a robust CD9 interactome with proximal proteins involved in numerous cellular pathways. Furthermore, CD9-mediated adherence required host cell surface receptors already present within the interactome but also recruitment of various proteins dependent on the infecting bacteria. Therefore, we present for the first time; i) a replete CD9 interactome within epithelial cells; ii) demonstrate the utility of CD9 as a universal organiser of ‘adhesion platforms’ for bacteria; iii) and provide new targets for investigation within bacterial adherence pathways and their downstream signalling responses.

## Conflicts of Interest

All other authors declare that the research was conducted in the absence of any commercial or financial relationships that could be construed as a potential conflict of interest.

## Author Contributions

Conceptualization, LRG and JGS; Methodology, LRG, JGS, MC, PAW and IFP; Formal Analysis, LRG, PAW and IFP; Investigation, LRG, PAW and IFP; Writing – Original Draft, LRG and IFP; Writing – Review & Editing, LRG, PAW, IFP, JGS and MC; Visualization, LRG; Funding Acquisition, LRG; Supervision, LRG and JGS.

## Acknowledgements

This study was funded by grants awarded to LRG from The Humane Research Trust (173280 to support PAW; https://www.humaneresearch.org.uk/), The Royal Society (RGS\R1\221334; https://www.royalsociety.org/), The Academy of Medical Sciences (SBF009\1010 to support IFP; https://acmedsci.ac.uk/) and and to MOC from the MRC (MR/X012220/1 to acquire MS instrumentation; https://www.ukri.org/councils/mrc/). The funders had no role in study design, data collection and analysis, decision to publish, or preparation of the manuscript. For the purpose of open access, the author has applied a Creative Commons Attribution (CC BY) licence to any Author Accepted Manuscript version arising.

The authors would like to acknowledge the biOMICS Mass Spectrometry Facility at The University of Sheffield for their help with mass spectrometry analysis.

## Supplementary Figures

**Supplementary Figure 1. CD9 removal affects meningococcal receptor expression.** (A) CD9 knockout has no effect on the number of cells expressing critical meningococcal receptors. % expressing cells of meningococcal receptors was determined by flow cytometry. WT or CD9^-/-^ cells were treated with an anti-CD9 antibody (602.29), an anti-CD46 antibody, an anti-CD147 antibody. A mouse IgG isotype control (JC1) was included and expression was determined using a FITC-conjugated secondary antibody. % change was calculated by comparing WT to CD9^-/-^ cells. (B) CEACAM expression was reduced upon CD9 knockout. Expression of CEACAM1 and CEACAM6 was determined by Western blot. Representative blot demonstrating expression of CEACAM1 and CEACAM6 in WT and CD9^-/-^ cells. Whole cell lysates of WT and CD9^-/-^ cells were electrophoresed and blotted. Blots were probed with anti-CEACAM1 or anti-CEACAM6 antibody. An anti-GAPDH antibody was used as a loading control. Densitometry was calculated through ImageJ analysis, removing background with an empty lane and normalising to the loading control. n≥1, mean ± SEM.

## Notes

### Competing Interest Statement

The authors have declared no competing interest.

