## Supplementary figures and images for "Dynamics of the CD9 interactome during bacterial infection of epithelial cells by proximity labelling proteomics"

### Supplementary Figure 1

#### Slide 1
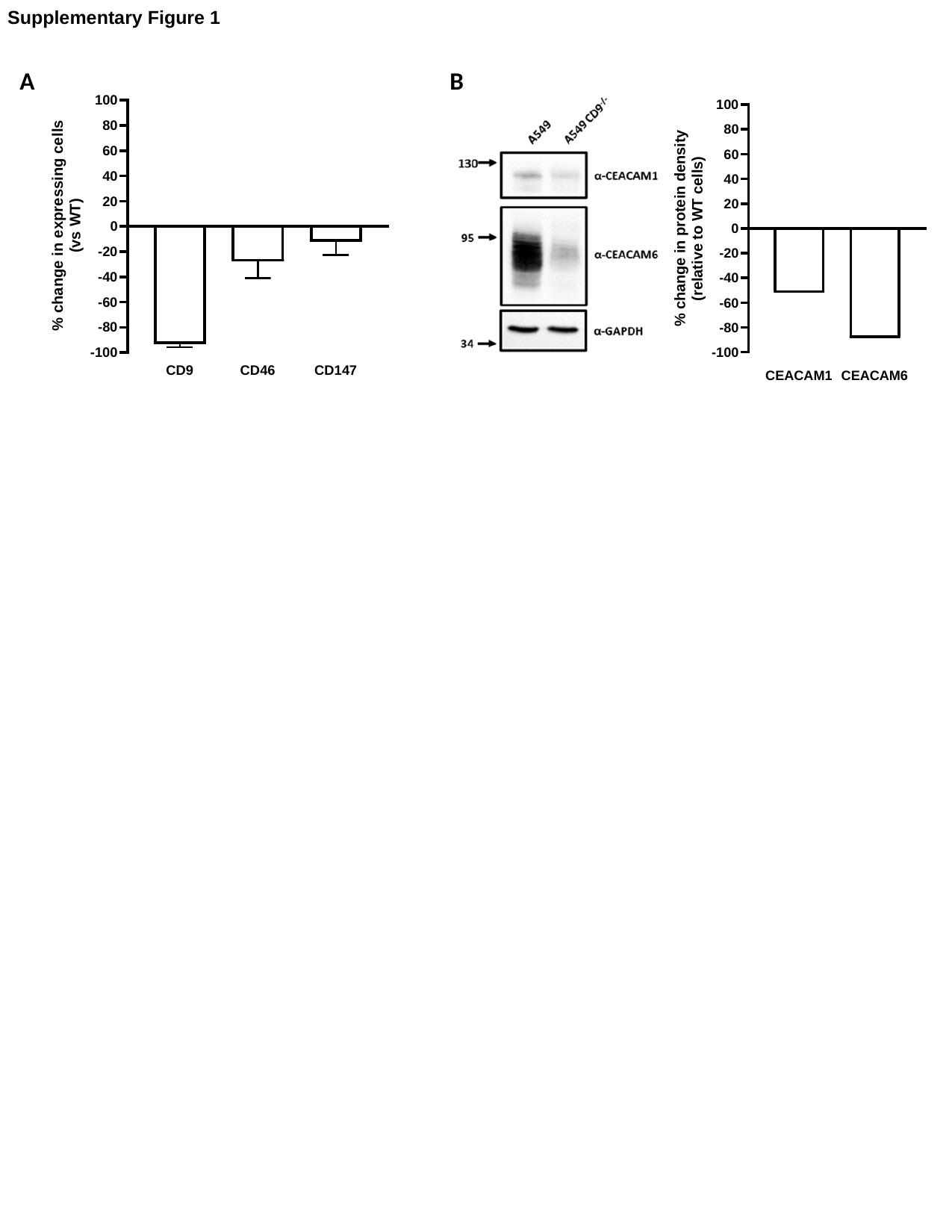

Supplementary Figure 1
A
B
